# Application of the thermal death time model in predicting thermal damage accumulation in plants

**DOI:** 10.1101/2024.01.29.577815

**Authors:** Andreas H Faber, Michael Ørsted, Bodil Kirstine Ehlers

## Abstract

The thermal death time (TDT) model suggests that the duration an organism can tolerate thermal stress decreases exponentially as the intensity of the temperature becomes more extreme. This model has been used to predict damage accumulation in ectotherm animals and plants under fluctuating thermal conditions. However, the critical assumption of the TDT model, which is additive damage accumulation, remains unverified for plants.
We assessed thermal damage in *Thymus vulgaris* under different heat and cold treatments and used TDT models to predict time to thermal failure of PSII. Additionally, thermal tolerance estimates from previous studies were used to create TDT models to assess the applicability of this framework in plants.
We show that thermal damage obtained at different stress intensities and durations is additive for both heat and cold stress, and that the TDT model can predict damage accumulation at both temperature extremes. Data from previous studies indicate a broad applicability of this approach across species, traits, and environments.
The TDT framework reveals a thermal tolerance landscape describing the exponential relationship between exposure duration, stress intensity and damage accumulation in plants. This thermal sensitivity emphasizes the potential impact of future thermal extremes on the mortality and distribution of plant species.

**Highlight:** This study highlights the applicability of the thermal death time model to plants, unveiling a distinct thermal tolerance landscape, extending across species and traits for assessing thermal stress impacts.

## Introduction

Temperature is one of the major factors determining the adaptation, natural selection and distribution of terrestrial plants (Gutschick & BassiriRad, 2003; Walther et al., 2005; Allen et al., 2010; Janská et al., 2010; Bertrand et al., 2011; Cramer et al., 2011; Lenoir, 2015; Nievola et al., 2017; Lancaster & Humphreys, 2020), and as the rate of climate change accelerates, the frequency and severity of extreme hot and cold temperature events increase around the globe (Easterling et al., 2000; IPCC, 2021; Rahmstorf & Coumou, 2011; Seneviratne et al., 2014). Extreme temperatures can impair physiological functions, growth, reproduction and consequently determine survival of plants by profoundly changing the structure and fluidity of cellular membranes, altering enzyme functions, and destroying proteins (Osmond et al., 1987; Sung et al., 2003; Hatfield & Prueger, 2015). Hence, understanding how species respond to these changes has never been more important. However, to understand how different species will respond to changing temperatures it is necessary to accurately determine how thermal tolerance and performance of plants is regulated through space and time.

Thermal tolerance in plants is often assessed through controlled ramping or static assays and is usually estimated in relation to stress intensity (e.g. upper and lower thermal tolerance limits and Thermal Performance Curves (TPCs)) (O’sullivan et al., 2013; Curtis et al., 2014; O’sullivan et al., 2017; Zhu et al., 2018; Lancaster & Humphreys, 2020; Sentinella et al., 2020; Geange et al., 2021; Wooliver et al., 2022). Across studies, stress durations can vary from anywhere between minutes to weeks and is sometimes considered negligible or simply dictated by the chosen stress assay type. However, thermal stress is a function of both stress intensity and duration, making it difficult to compare across studies and/or relate estimates to field conditions where plants may experience large fluctuations in both intensity and duration of temperature exposure. For example, leaf temperatures in the field can rise dramatically within minutes and can exceed air temperatures of more than 10°C (Wise et al., 2004; Vogel, 2005; Rey-Sanchez et al., 2016; Fauset et al., 2019), which is enough to significantly suppress photosynthetic activity and lead to oxidative stress (Vallélian-Bindschelder et al., 1998). Not only is it difficult to relate lab derived thermal tolerance estimates to such conditions, but it is also difficult to identify and characterize different thermal stress-conditions such as those that are suboptimal, induce protective mechanisms or result in acute damage accumulation when the stress varies greatly in both intensity and duration.

In the literature on thermal tolerance of ectotherm animals, a promising solution to reconcile several of these problems has been proposed in the recently (re)introduced Thermal Death Time (TDT) model which combines information on the severity and duration of stressful temperature exposure (Rezende et al., 2014; Jørgensen et al., 2019; Jørgensen et al., 2021). The TDT model is relevant in the stressful temperature range where an organism may experience acute thermal damage accumulation, and the model can predict responses to fluctuating temperature conditions such as in the field. The underlying theory behind this approach is that there exists an exponential relationship between the intensity of the stress and the amount of time an organism can tolerate before it succumbs to thermal stress (Fig. **1A**). This association has been known for more than a century in ectothermic organisms (Bigelow 1921; Fry et al., 1946; Maynard, 1957; Kilgour & McCauley, 1986) and plants (Collander, 1924; Brown & Crozier, 1927; Berkley & Berkley, 1933; Sapper, 1935; Lorenz, 1939; Alexandrov 1964). The TDT curve is traditionally expressed as a linear regression between temperature and log_10_ transformed exposure durations (Fig. **1B**). Thermal damage described in the TDT model can occur at the population, individual or cellular level and the model should apply for both heat and cold stress. TDT curves assume that thermal tolerance is embedded in a three-dimensional space known as a Thermal Tolerance Landscape (TTL) (Fig. **1C**). A TTL describes the temperature–time interaction of tolerance, along with information of percentwise thermal damage accumulation (Jørgensen et al., 2021) or percentwise population mortality level (Rezende et al., 2014). Thus, for each damage level (for example 50%) there is a specific TDT curve, and multiple TDT curves jointly form the TTL for a population (Rezende et al., 2014). If the TDT curve framework is true, an organism may move around in this TTL and will succumb to thermal stress when 100% damage has been received, regardless of whether it was received at one single, or several different temperatures. This implies that damage accumulation received at different temperatures is additive (Fry, 1947, 1971; Fry et al., 1946; Kashmeery & Bowler, 1977; Jørgensen et al., 2021). Additive damage accumulations have been demonstrated for ectotherm animals (Fry et al., 1946; Jørgensen et al., 2021), however, the assumption of additivity has never been tested in plants. If damage accumulates additively, it should be possible to use TDT model parameters derived from static experiments to accurately predict the time when an organism succumbs to thermal stress during 1) combinations of different static exposures, 2) gradually increasing or decreasing temperatures where a linear temperature increase results in an exponentially increasing rate of thermal failure and/or 3) fluctuating temperature exposures (Ørsted et al., 2022). This could facilitate the comparison of thermal tolerance and assessment of thermal conditions that lead to damage accumulation across different studies and methodologies. Consequently, it would provide a unified approach to evaluate the impacts of thermal stress on plants.

**Fig. 1.**
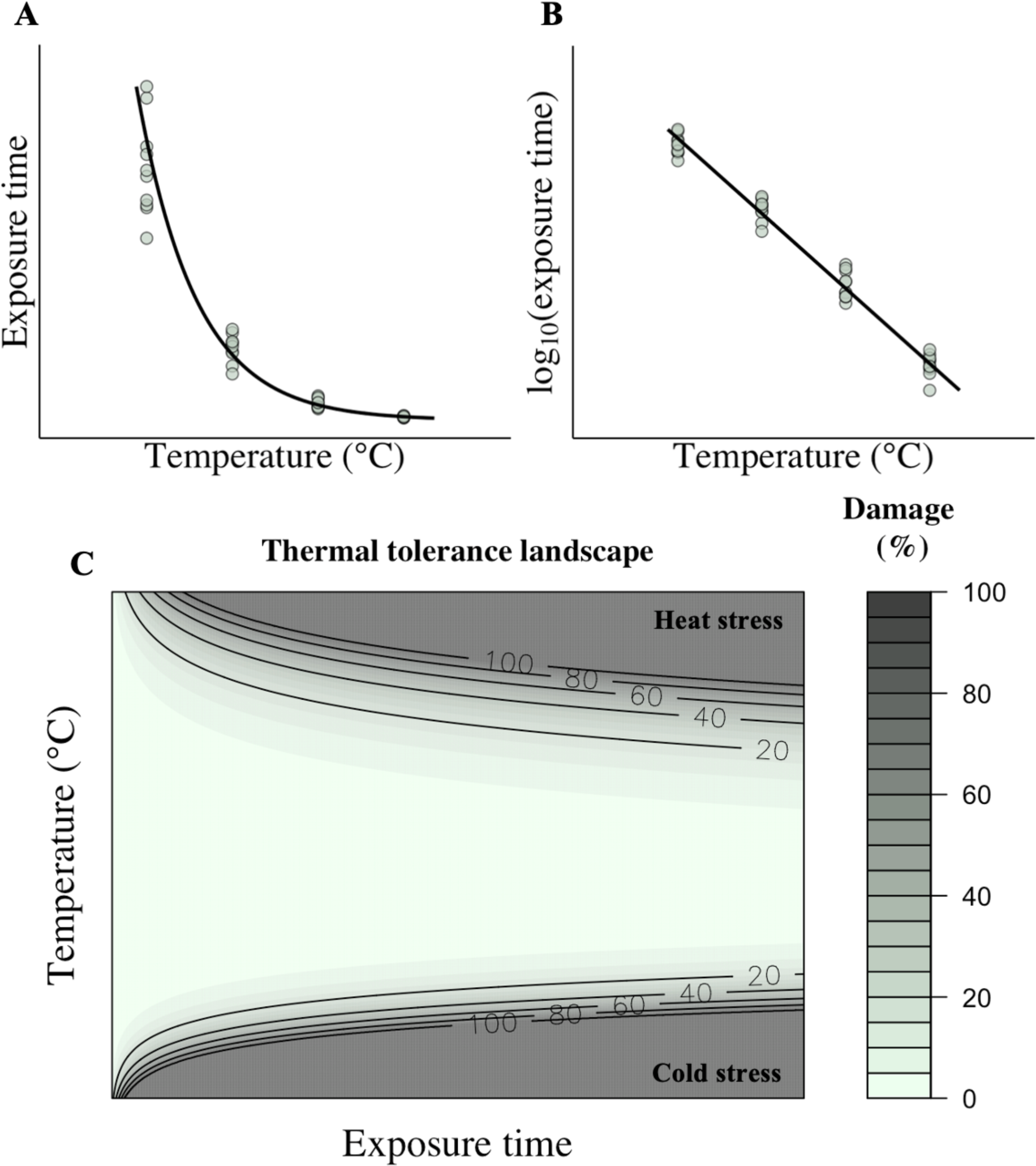
Conceptual figures of **A)** the exponential relationship with thermal tolerance (e.g. time of damage or death) and stress temperature (for heat stress) and, **B)** the same data shown as a thermal death time (TDT) curve where thermal tolerance is expressed as log_10t_ = ß*stress temperature + α where ß and α is the slope and intercept, respectively. **C)** Thermal tolerance landscape calculated using TDT curves of heat and cold thermal tolerance response where damage accumulation is modeled as a function of intensity and exposure time of stress.

Only very recently has it been demonstrated how TDT curves derived from static assays can be used to predict plant mortality under field conditions (Cook et al., 2023, Preprint; Neuner & Buchner, 2023). These studies highlight a promising research agenda in plant ecophysiology emphasizing the need for further investigation in this direction. However, the extent to which the TDT framework applies across plant species, traits, and environmental conditions remains largely unexplored. This is especially true for cold stress, as only few studies have specifically investigated this relationship (Levitt, 1980), while others have indirectly shown its relevance (Bigras et al., 2004). To be broadly applicable, the TDT framework requires damage to exhibit a cumulative nature. This implies that damage resulting from a single instance of stress accumulates and contributes to the overall damage, arising from a combination of stress intensities and exposure durations. Therefore, testing this additivity for both heat and cold stress is essential before TDT curves can be reliably employed for predictive purposes. Since plants share many basic characteristics of the biological processes responsible for thermal failure in ectotherms (e.g. protein denaturation, inactivation of enzymes and membrane dysfunction) (Levitt 1980; Schmidt-Nielsen, 1997; Nievola et al., 2017) we hypothesize that thermal damage accumulation is additive for plants as well.

The aim of this study was firstly: to assess whether damage accumulation exhibits the TDT relationship in response to both heat and cold stress, as well as to examine the assumption of additive damage accumulation in plants under both types of stress. Secondly, we sought to evaluate the applicability of the TDT framework in plants across different species and traits (stress traits). Using the Mediterranean plant Thyme (*Thymus vulgaris* L.), we tested whether thermal damage accumulation is additive for both heat and cold stress. To do so, we experimentally constructed TDT curves for heat and cold stress. We then used these curves to predict the time to thermal failure under different consecutive temperature stress treatments, and subsequently tested the accuracy of these predictions. The plant trait we used to test thermal damage was maximum quantum efficiency of photosystem two (PSII). Given the high sensitivity of plant photosynthesis to heat (Berry et al., 1980; Havaux et al., 1991; Georgieva & Yordanov, 1994) and cold stress (Huner et al., 1993; Allen & Ort, 2001; Devacht et al., 2009; Mai et al., 2009; Chaves et al., 2015), particularly with respect to damage to PSII (Fracheboud et al., 1999; Georgieva & Lichtenthaler, 2006; Mathur et al., 2014; Chaves et al., 2015), we employed damage to PSII as an indicator of thermal failure. This choice was based on its ease of measurement, its established reflection of tissue-level damage (Bilger et al., 1984), and its prior use and suitability for predicting mortality of plants in the field (Cook et al., 2023, Preprint; Neuner & Buchner, 2023). In addition to our experimental approach, we also searched the literature for studies on plant thermal tolerance using different plant traits. We subtracted the data from these studies to construct TDT model across different species, traits, methods, and environmental conditions in order to determine the broad applicability of this approach.

## Materials and methods

### Growth conditions

Thyme seeds were collected in June 2022 in Southern France from natural populations growing in the garrigue area in the mountains of “la Séranne” (43,88N, 3,67E) and stored in paper bags at room temperature until use. The plants were grown in Denmark in August 2022 in the greenhouse at Dept. of Ecoscience, Aarhus University (56.198°N, 10.155°E). Seeds were grown in trays filled with a 5 cm layer of Mediterranean peat- based soil and the trays were kept humid under a white plastic sheet until germination. The seedlings were transferred to individual 15 cm pots when they reached ∼10 cm in height in October 2022 and placed on growing benches. The CO_2_ concentration was set at ambient conditions (∼410 ppm) and the temperature ranged between 16 and 26 °C with a relative humidity of ∼60%. During the day, the natural sunlight was supplemented by LED lamps (FL300 Grow, Senmatic, Søndersø, Denmark) when the photosynthetic photon flux density (PPFD) was below 150 µmol m^-2^ s^-1^. The climate parameters were controlled by a climate control unit (LCC4, Senmatic, Søndersø, Denmark). Plants were irrigated one to two times a week and supplied with a NPK nutrition solution.

### Sampling and static assay protocol

In this study, thermal tolerance was evaluated by assessing the efficiency of PSII following thermal stress. The sensitivity of PSII to thermal stress was determined using chlorophyll fluorometry to measure the maximum quantum efficiency of PSII or *F*_V_/*F*_M_. *F*_V_ and *F*_M_ are the variable and maximum fluorescence of dark-adapted leaves, respectively, where *F*_V_ is calculated as the difference between dark-adapted maximum and minimum fluorescence (*F*_o_). *F*_V_/*F*_M_ represents the capacity of PSII to convert absorbed light into photochemical energy (Berry & Bjorkman, 1980; Seemann et al., 1984; Willits & Peet, 2001; Knight & Ackerly, 2002, 2003; Baker & Rosenquist, 2004). A decrease in *F*_V_/*F*_M_ of more than ∼50% indicates dysfunction in the photosynthetic machinery (Schreiber & Berry, 1977; Downton & Berry, 1982; Knight & Ackerly, 2003; Sastry & Barua, 2017), leading to visual tissue level damage (Bilger et al., 1984).

To generate TDT curves, heat and cold stress tolerance was estimated for a range of constant temperatures and exposure durations. Four batches of samples were collected for each temperature treatment and a batch consisted of 10 samples for heat and 20 samples for cold treatments. The samples were obtained from the outer tips of branches, approximately 2 cm in length. 10 different plants were sampled for heat treatments, while eight plants were sampled for cold treatments. The samples were collected at 8-10 AM and only branches with fully expanded and healthy leaves of similar age were randomly sampled from each plant. All plants were represented by minimum one sample in each batch.

Following collection, the samples in each batch were placed in 2 ml micro centrifuge tubes and kept in darkness at room temperature (20-22°C) for 1-3 hours. Subsequently, the maximum quantum efficiency of PSII before stress exposure was estimated by measuring *F*_V_/*F*_M_ using a chlorophyll fluorometer (FlourPen FP110/S; Photon Systems Instruments, Drásov, Czech Republic). Next, the batches were placed under actinic light at ∼175 µmol photons m^-2^ s^-1^ for 30 minutes and submerged in water baths (accuracy of 0.02 °C, LAUDA ECO RE 1050, LAUDA, NJ, USA), to minimise the temperature difference between plant material and treatment temperature. Each batch was individually withdrawn from the water bath after a specific treatment duration. For the heat stress treatments, the temperatures were fixed at 43, 43.5, 44, 44.5, 45.5, 46, 47 or 47.5 °C. Each batch was removed from the treatment following different exposure times which ranged from 15 and 480 minutes. The variation in exposure times was chosen with respect to the stress intensity, with lower stress intensities requiring longer exposure times. This was implemented to achieve the desired stress levels for the different temperature treatments. During heat stress treatments the samples were exposed ∼95 µmol photons m^-2^ s^-1^ since heat stress is usually accompanied by light under natural conditions and allows for the onset of light-dependent reactions involved in thermal protection and recovery (Curtis et al., 2014). The light intensity was measured at the level of the samples in the water bath with the FlourPen concealed in a transparent plastic bag. For the cold stress treatments, the temperatures were fixed at -8, -7.5, -7 or -6.5 °C. Each batch was removed from the treatment following different exposure times which ranged from 5 to 2540 minutes. The samples were not exposed to light during the cold stress treatments, as a cooling liquid was used to decrease the water temperature below freezing which did not allow light penetration. Following the stress treatments, the batches were exposed to ∼175 µmol photons m^-2^ s^-1^ for 60 minutes and subsequently dark adapted for 16- 19 hours, before *F*_V_/*F*_M_ was measured again to estimate the consequential accumulated damage to PSII. This process was conducted to estimate the complete dose of damage incurred at a specific stress temperature, a process that may accumulate many hours following the thermal stress (Curtis et al., 2014). One to three temperature treatments were conducted per day, and the same water bath was used for all batches for each individual temperature treatment. For each day of conducting experiments, *F*_V_/*F*_M_ was measured on 10 control samples in parallel to the heat and cold stress treatments. The control samples were kept at constant room temperature next to the water baths and was given the same dark/light treatments as the heat and cold stress treatments. The control samples were used to confirm that any decrease in *F*_V_/*F*_M_ could be attributed to thermal stress and showed that *F*_V_/*F*_M_ was identical before and after the experimental period when the samples were not stressed (Fig. **S1A-B**).

### Additive damage accumulation assay

To assess whether thermal damage accumulates in an additive manner for both heat and cold stress, a set of experiments was conducted in which batches of samples were exposed to two consecutive temperature intensities. The same light treatments as in the static assays was applied for these experiments and *F*_V_/*F*_M_ was measured before and after the thermal stress to estimate the resulting damage to PSII. Similarly, *F*_V_/*F*_M_ was measured on 10 control samples each day of measuring in parallel to the stress treatments (Fig. **S1C-D**). For the heat stress experiments, four batches, consisting of 10 samples, were initially stressed in one water bath at 44°C for 30 minutes. Subsequently, they were moved to another water bath at either 44.5, 45, 45.5, 46, 46.5 or 47°C. Each batch was removed from the treatment following different exposure times, ranging from 15 to 135 minutes. For the cold stress experiments, four batches consisting of 20 samples, were first stressed in a water bath at -6.5°C for five hours. They were then moved to another water bath at either -7, -7.5 or -8 °C, where each batch was removed at different exposure times ranging from 5 to 360 minutes. One to four temperature treatments were conducted per day, and the same water bath was used for all batches for each individual temperature treatment.

### Thermal death time curves in previous studies

To evaluate the general applicability of the TDT framework to plants, we examined thermal tolerance estimates from previous studies. The aim of this analysis was not to provide a full comprehensive overview of all studies showing the TDT relationship, but rather to determine if the TDT framework can be applied for a range of species and traits based on various methodologies and environmental conditions. To provide illustrative examples, we incorporated data from Alexandrov (1964), where TDT curves for heat stress were established based on assays of protoplasmic streaming, and data from Bělehrádek & Melichar, (1930), where TDT curves for heat stress were based on live tissue staining with Ruzicka’s neutral rose-methylene blue. Furthermore, the investigation aimed to assess the feasibility of creating TDT curves using data from studies that presented thermal tolerance as a function of stress intensity and exposure duration, rather than explicitly providing TDT curves. In this context, we provided illustrative examples through generating TDT curves by log_10_ transforming time to thermal failure based on a 50% reduction in seed viability of heat stress from Dahlquist et al., (2007) and on browning of twigs due to cold injury from Sakai, (1956), as a function of the intensity of the stress.

### Statistical analysis

Generalized additive models (GAMs) were used to estimate the decrease in *F*_V_/*F*_M_ following the stress treatments and determine the critical temperature at which a lethal dose of damage to PSII occurs following a specific time of exposure (CT_!_). For heat stress, GAMs were fitted to *F*_V_/*F*_M_ measurements as a function of stress duration, with CT_!_ indicating the time when *F*_V_/*F*_M_ decreased by 50%. In contrast, for cold stress, where *F*_V_/*F*_M_ either remained largely unaffected or very close to zero, GAMs were fitted to the percentage of individual samples within each batch that did not exhibit more than a 50% decrease in *F*_V_/*F*_M_ as a function of stress duration. In this case, CT_!_was estimated as the time when 50% of the samples had more than a 50% decrease in *F*_V_/*F*_M_.

The GAMs were fitted using automated smoothness selection in the *mgcv* library in R v.4.1.0 (R Core Team, 2021) and restricted maximum likelihood (REML) (Wood, 2017). The GAMs had the following components:

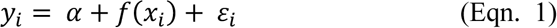

Where *y_i_* is the observation at time *x_i_*, α is the intercept, *f*(*x*_*i*_) is a smooth function and ε_*i*_ is the residual error. This approach is non-parametric and makes no *a priori* assumption about the functional relationship between variables (Wood, 2017), allowing the depiction of the empirical trend of the response over time without restrictions. The only exception was for the fixed temperature treatment at -8°C where it was necessary to apply a log transformation of *x_i_* to fit the model. For the static assays used to generate TDT curves, the GAMs were fitted by making 10.000 random draws from the multivariate normal distribution of the model parameters. For each static temperature, 95% of the random draws closets to the mean was used to estimate CT_!_. For the additive damage accumulation experiments, the GAMs were fitted with 95% pointwise confidence intervals and the mean of the fitted GAMs was used to estimate CT_!_. Before analysis, model assumptions of normality and homoscedasticity of residuals were assessed and verified.

TDT curves were generated by fitting a linear regression to the log_10_ transformed time values representing the CT_!_random draws estimated during the static experiments, and these time values were plotted against the corresponding temperature intensities. The TDT curves were described as:

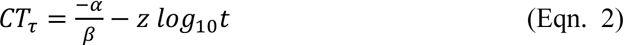

Where CT_τ_ is the critical temperature at which a lethal dose of damage occurs following a specific time of exposure (°C) (i.e. a 50% reduction in *F*_V_/*F*_M_ for heat stress or a reduction of more than 50% in *F*_V_/*F*_M_ in 50% of the samples for cold stress), α and β is the intercept and slope of the linear regression model, respectively, *t* is the elapsed exposure time, and *z* is a constant that characterizes the sensitivity to temperature change (i.e. the temperature change resulting in one order of magnitude change in time to heat failure). The parameter *z* can be calculated by the slope of the TDT curve as *z* = -1/slope and is analogous to Q_10_, which denotes the factor by which thermal tolerance change for every 10 °C change in temperature, where Q_10_ = 10^10/*z*^. Across the TDT curves of Thyme and the species found in the literature, CT_!)(*+,_ or CT_!)-./0_ was calculated using equation two, which represents the critical temperature required to inflict a lethal dose of damage following 30 minutes or 35 days of stress exposure, respectively. In addition, the thermal sensitivity was expressed as a Q_10_ for all species.

For testing whether thermal damage accumulation is additive for Thyme, CT_!_ was predicted for the additive damage accumulation assays with two consecutive temperature intensities based on the derived TDT model parameters according to Jørgensen et al., 2021:

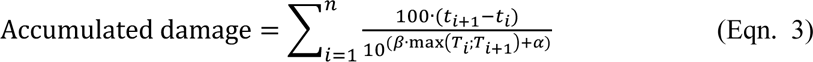

Where the accumulated damage is calculated over each time interval *i* unit t_e_ which is the time interval for which the total accumulated damage is calculated. The β and α are the slope and intercept of the TDT curve, respectively. The denominator is the fraction of the tolerable exposure duration for the maximum temperature (of T_i_ and T_i+1_) in each time interval. For heat and cold stress, 100% damage accumulation is equal to CT_!_. An example of how to predict the time where an organism succumbs to thermal stress when the temperature fluctuates is demonstrated in the supplementary R script (**Notes S1**). The predicted and observed CT_!_ values were compared using a one-sample two-sided t-test to estimate whether the TDT curves reliably predicted the onset of a lethal dose of damage for both heat and cold stress.

## Results

### Damage accumulates exponentially with the intensity of heat and cold stress

There was a time dependent accumulation of damage to PSII across all the fixed temperature treatments for both heat and cold stress (Fig. **S2, S3**). Similar to an exponential decay, *F*_V_/*F*_M_ gradually decreased with the duration of the heat stress (Fig. **S2**). For cold stress, a similar decrease was observed for the percentage of individual samples where *F*_V_/*F*_M_ was above 50% (Fig. **S3**). This time dependent accumulation of damage yielded CT_!_ estimates for both types of stress, which represents the critical temperature at which a lethal dose of damage occurs following a specific time of exposure.

The exposure time for CT_!_ to occur for heat and cold stress exhibited a linear relationship with the stress temperature (Fig. **2A,C**; *R*^2^ = 0.99, *P* < 0.05 and *R*^2^ = 0.98, *P* < 0.05, for heat and cold stress, respectively), showing that the thermal tolerance decreased exponentially with the stress intensity for both types of stress. This linear relationship clearly demonstrates that TDT models can be extrapolated and used to predict damage when the stress is provided through a constant temperature assay. As illustrated by the TTLs generated by the TDT models, damage accumulation was relatively gradual for heat stress in comparison to cold stress (Fig. **2B,D**). The time for receiving 100% heat damage varied between a few minutes to around six hours for a 4.5°C change in temperature. For cold stress, this time varied between a few minutes to ∼30 hours for only a 1.5°C change in temperature. CT_τ30min,_ and the Q_10_ value for heat stress was predicted as 45.9 °C and 3,981, respectively. For cold stress, CT_τ30min,_ and the Q_10_ was predicted as -7.4 °C and 3.98*10^19^, respectively. These results underline the fact that the rate of damage accumulation differs by several orders of magnitude between the two stress types.

**Fig. 2.**
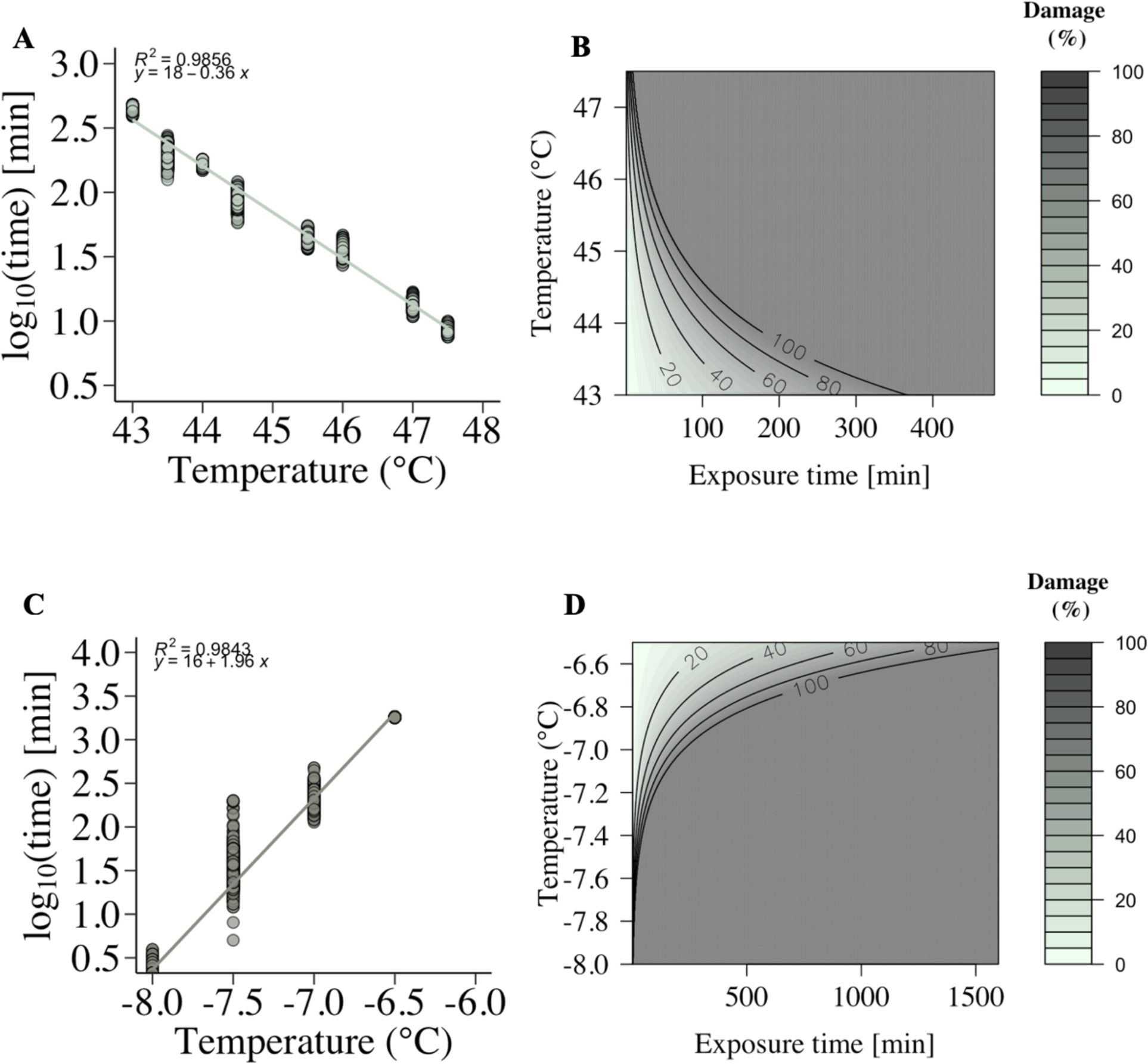
Thermal death time curves of **A)** heat and **C)** cold thermal tolerance measured at a range of constant temperature treatments for *Thymus vulgaris*. Both heat and cold thermal tolerance is represented as CT_!_ which denotes the critical temperature at which a lethal dose of damage to PSII occurs following a specific time of exposure. The y-axis depicts the log_10_ transformed time in minutes for CT_!_ to occur and the x-axis depicts the intensity of the thermal stress (°C). CT_!_was calculated for each temperature treatment using generalized additive models, and the individual points illustrates CT_!_ for 95% of 10.000 random draws closest to the mean from the multivariate normal distribution of the model parameters. The TDT curves were used to generate thermal tolerance landscapes of **B)** heat and **D)** cold stress. The thermal tolerance landscapes were calculated using the TDT curves of the heat and cold thermal tolerance response, where damage accumulation is modeled as a function of intensity and exposure time (minutes) of stress. For heat and cold thermal tolerance, 100% damage is equal to CT_!_.

### Damage accumulation from heat and cold stress is additive

To assess whether thermal damage accumulates in an additive manner, plant material was stressed by a series of two consecutive temperatures and TDT curve parameters were used to predict CT_!_ for each scenario. For heat stress, samples were subjected to initial stress at 44°C for 30 minutes. Subsequently, the samples were transferred to either 44.5 °C, 45 °C, 45.5 °C, 46 °C, 46.5 °C or 47 °C for different durations and the difference between the observed CT_!_ and the predicted was 12.35, 0.73, 11.34, 0.7, 12.74 and 5.12 minutes, respectively (Fig. **3A** and Fig. **S4**). In addition, the predicted CT_!_ was generally lower that the observed CT_!_ but not statistically different (*t* = -0.27881, *P* > 0.05). For cold stress, samples were initially stressed at -6.5 °C for five hours. Subsequently the samples were transferred to either -7 °C, -7 °C, -7.5, -7.5 °C, -8 °C or -8 °C for different durations and the difference between the observed and predicted CT_!_was 113.27, 35.7, 32.05, 6.06 and 4.38 minutes, respectively (Fig. **3B** and Fig. **S5**). The predictions of CT_!_ for cold stress was generally lower than the observed CT_!_ but not statistically different (*t* = -0.6013, *P* > 0.05) but in one case CT_!_ was not estimated in the experimental timescale (Fig. **S5B**).

**Fig. 3.**
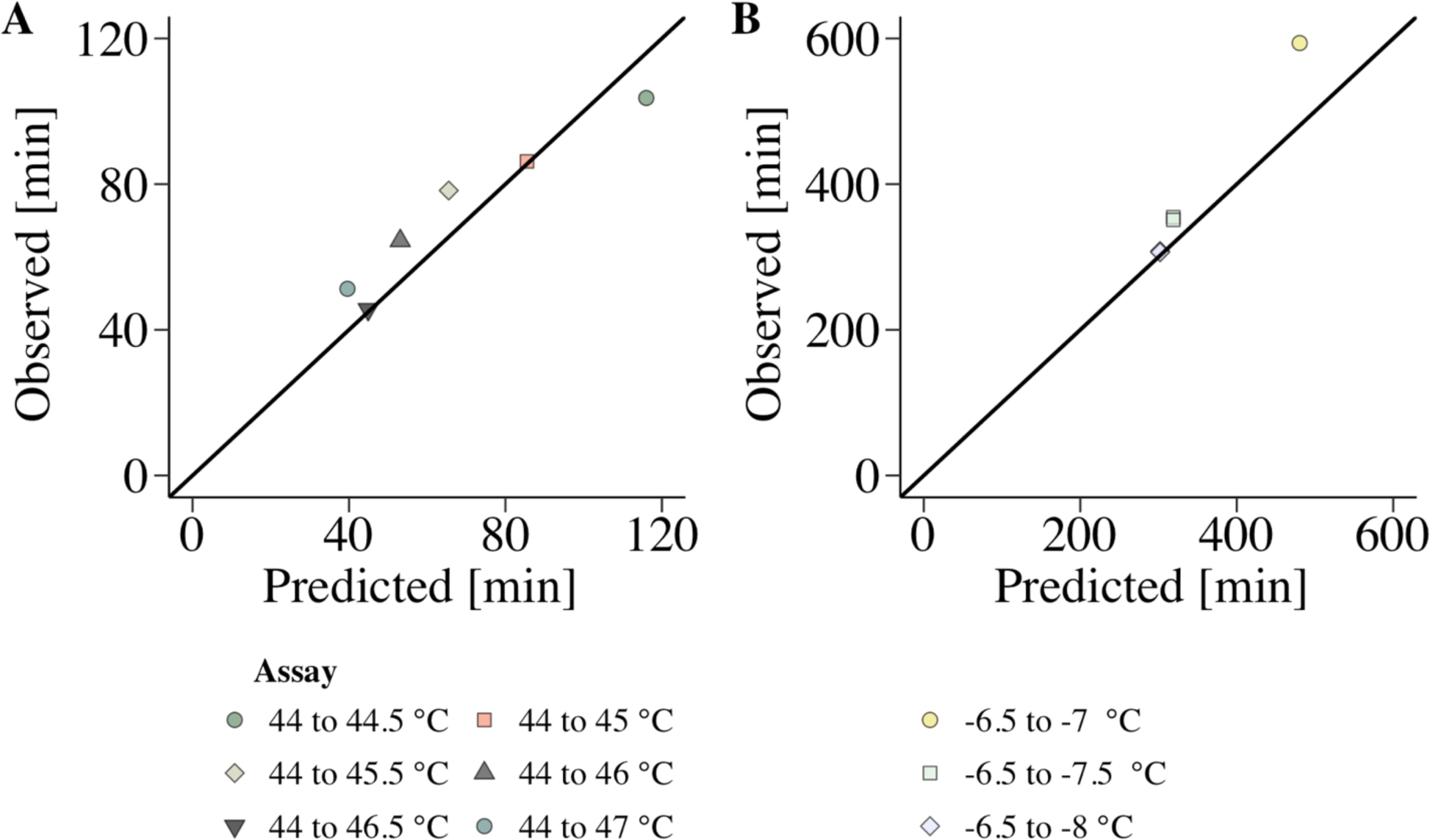
The difference in minutes between the observed and predicted CT_!_for **A)** heat and **B)** cold stress for *Thymus vulgaris*. For both heat and cold thermal tolerance CT_!_ denotes the critical temperature at which a lethal dose of damage to PSII occurs following a specific time of exposure. For heat stress, samples were stressed at 44 °C for 30 minutes and subsequently transferred to either 44.5 °C, 45 °C, 45.5 °C, 46 °C, 46.5 °C or 47 °C for different durations. For cold stress, samples were stressed at -6.5 °C for five hours and subsequently transferred to either -7 °C, -7.5 °C, or -8 °C for different durations. For cold stress, the experiment was conducted twice, but CT_!_ was not estimated in one of the treatments where the samples were exposed to -6.5 °C followed by -7 °C. The predicted CT_!_ was calculated using the TDT curves of the heat and cold thermal tolerance response, where damage accumulation is modeled as a function of intensity and exposure time (minutes) of stress. Different symbols indicate different stress treatments.

### The TDT model can be applied for a range of species and traits

Several studies have generated TDT curves on plants (Collander, 1924; Brown & Crozier, 1927; Bělehrádek & Melichar, 1930; Berkley & Berkley, 1933; Sapper, 1935; Lorenz, 1939; Alexandrov, 1956; Alexandrov & Fel’dman, 1958; Gorban’, 1961; Fel’dman & Liutova, 1962; Alexandrov, 1964; Wright, 1970; Nishiyama, 1975; Levitt, 1980; Colombo & Timmer, 1992; Hara, 2005; Peter et al., 2009; Cook, 2021; Cook et al., 2023, Preprint; Müller et al., 2023; Neuner & Buchner; 2023) (examples are shown in Fig. 4A and **C**). However, some studies have investigated thermal tolerance throughout time without explicitly representing it through TDT curves (Sakai, 1956; Dahlquist et al., 2007). In such cases, it was still feasible to generate TDT curves based on the provided thermal tolerance estimates (Fig. 4B and **D**). Collectively, these results suggests that TDT curves can be generated for different traits and species for heat stress, such as the cessation of protoplasmic streaming of leaf cells (Fig. **4A**), seed viability (Fig. **4B**) and staining of live tissues (Fig. **4C**) as well as browning of tissues following freezing during cold stress (Fig. **4D**). The TDT curves illustrate that thermal tolerance estimates can be compared not only by Q_10_ but also by the amount of homeostatic failure that is applied. For example, the Q_10_ values of the TDT models in Fig. **4A** are similar across different species, but the species differ greatly in thermal tolerance when comparing the temperature resulting in 100% damage following a 30-minute exposure duration (CT_!)(*+,_). Although confounded by effects of species variation, the Q_10_ values and CT_!)(*+,_ seems to vary across different traits (Fig. **4A-C**) where the Q_10_ for cold stress is substantially lower compared to heat stress (Fig. **4D**). This much slower accumulation of damage is reflected by the fact that the species exposed to cold stress had to be compared in stress durations of days instead of minutes (CT_!)-./0_) (Fig. **4D**). Interestingly, Fig. **4C** depicts a high temperature range which is separated from a lower temperature range with a sudden break from linearity.

**Fig. 4.**
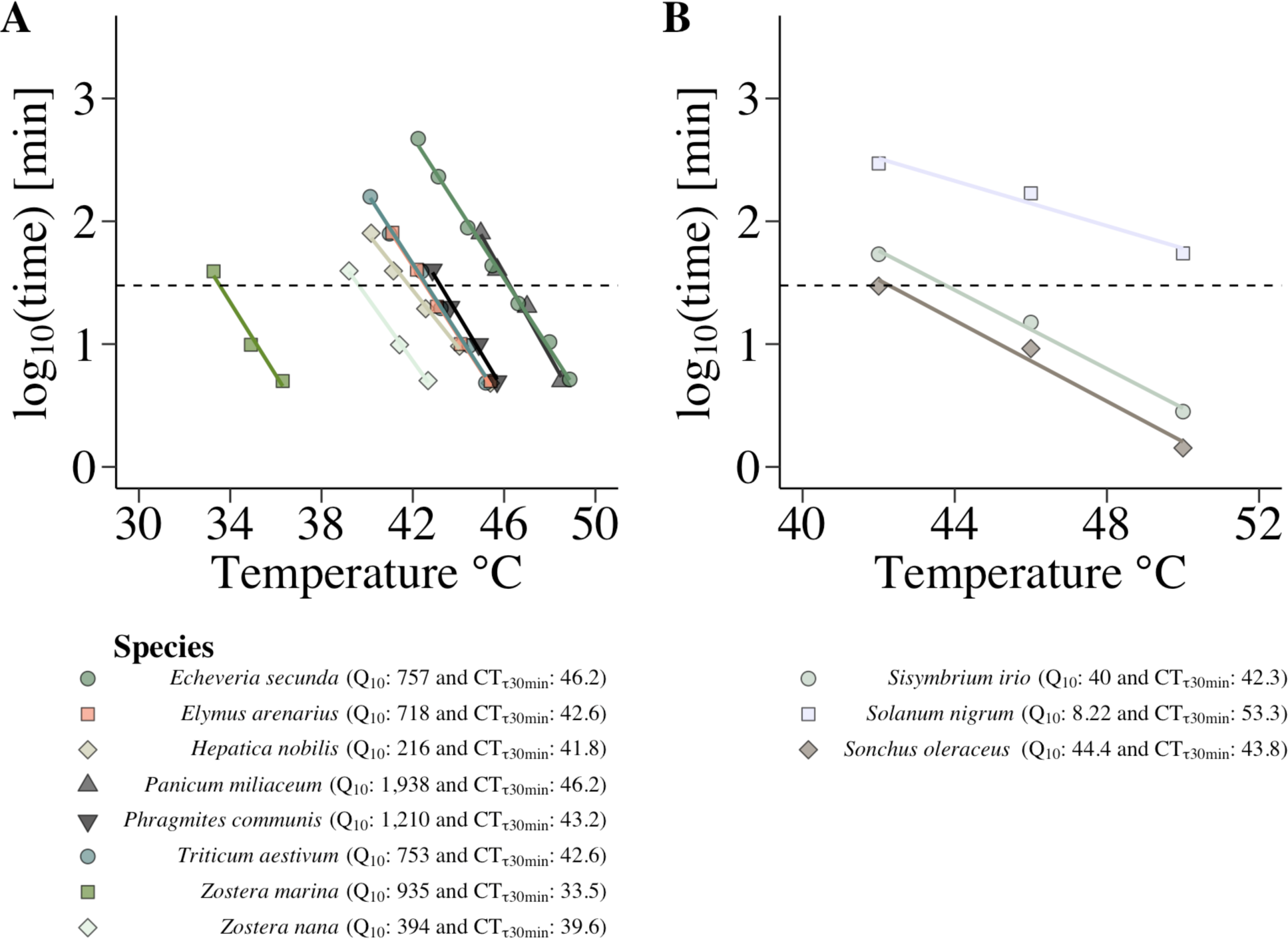
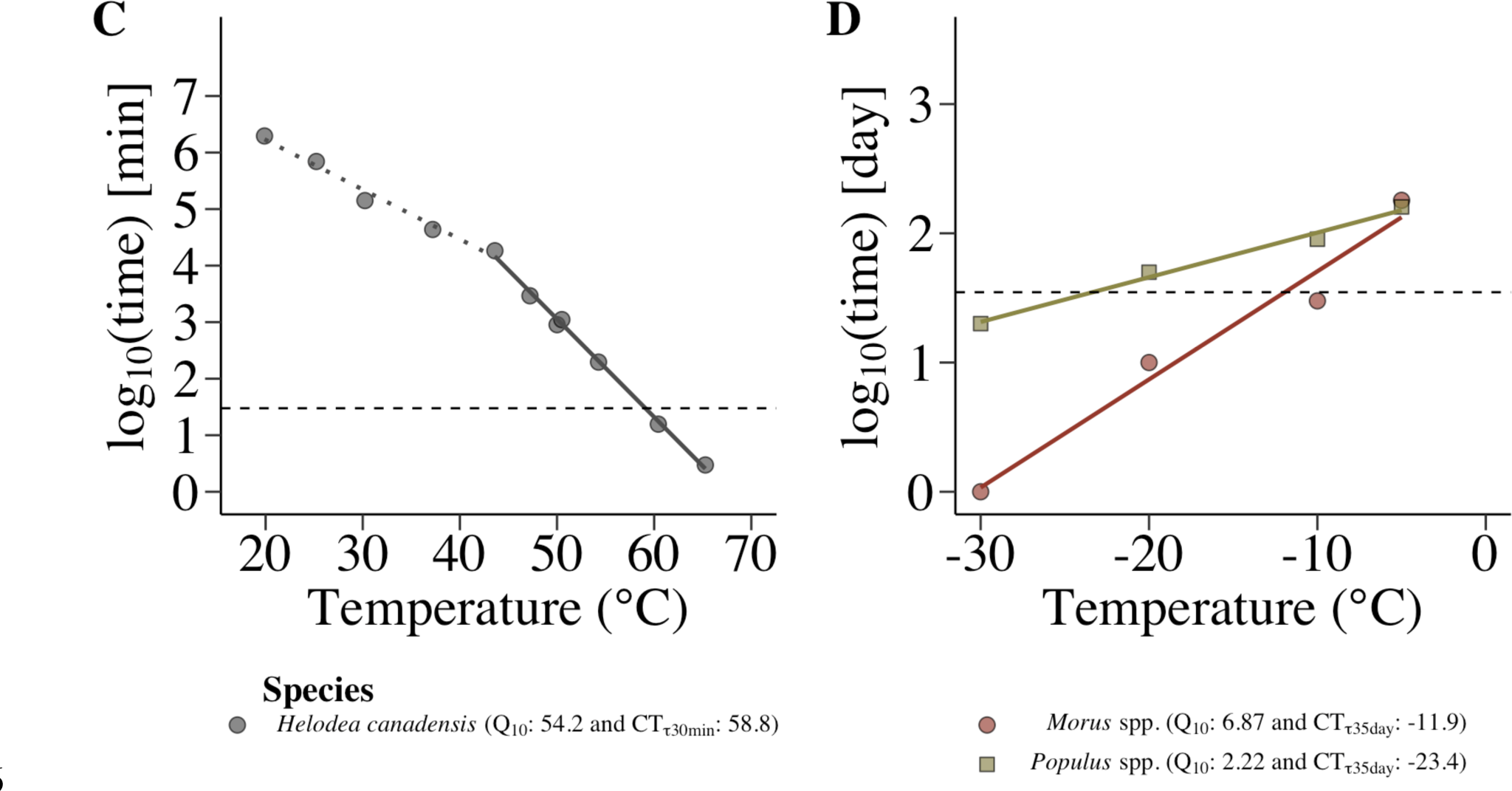
TDT models based on literature data of **A)** cessation of protoplasmic streaming of leaf cells of *Echeveria secunda* Booth. (data from Alexandrov, (1964)), *Elymus arenarius* L. (data from Alexandrov, (1956)), *Hepatica nobilis* Schreb. (data from Alexandrov, (1964)), *Panicum miliaceum* L. (data from Alexandrov, (1956)), *Phragmites communis* Trin. (data from Alexandrov, (1956)), *Triticum aestivum* (data from Alexandrov, (1956)), *Zostera marina* L. (data from Fel’dman & Liutova, (1962)) and *Zostera nana* Roth. (data from Fel’dman & Liutova, (1962)) following heat stress, **B)** 50% seed viability of *Sisymbrium irio* L. (data from Dahlquist et al., (2007)), *Solanum nigrum* L. (data from Dahlquist et al., (2007)) and *Sonchus oleraceus* L. (data from Dahlquist et al., (2007)) following heat stress, and **c)** live tissue staining with Ruzicka’s neutral rose-methylene blue of *Helodea canadensis* Michx. following heat stress depicting a high temperature range which is separated from a lower temperature range with a sudden break from linearity (data from Bělehrádek & Melichar, (1930)) and **D)** survival determined by browning of twigs of *Morus* spp. L. (data from Sakai, (1956)) and *Populus* spp. L. (data from Sakai, (1956)) following freezing during cold stress. The y-axis depicts log_10_ transformed time in minutes (**A-C**) and days (**D**), and the dotted horizontal lines denotes a 30-minute (**A-C**) and a 35-day (**D**) exposure time.

## Discussion

### The TDT model reveals two distinct temperature ranges

This study demonstrates that thermal damage accumulation to PSII is additive for a specific range of temperatures for both heat and cold stress in Thyme. Importantly, in the temperature range where damage is additive, the fundamental condition for this additivity is the absence of recovery mechanisms. This condition aligns with the underlying principle of the TDT curve, where time to reach a given level of damage is dictated by the intensity of the stress. This result underlines the potential for accurately predicting thermal damage accumulation in plants with the TDT framework, even under varying temperature and exposure duration conditions, supporting the recent approaches of estimating mortality of plants under field conditions (Cook et al., 2023, Preprint; Neuner & Buchner, 2023). Furthermore, data from previous studies show that TDT curves can be generated on a range of different plant species, traits, and environmental conditions. This diversity in traits that exhibit the TDT relationship indicates a broad applicability of the TDT framework. To date it has only been demonstrated that thermal damage accumulation is additive for the brook trout (*Salvelinus fontinalis* Mitchchill) and vinegar fly (*Drosophila melanogaster* Meigen) for heat stress (Fry et al., 1946; Jørgensen et al., 2021) and now for Thyme for both heat and cold stress. This additive damage accumulation may only occur when the stress intensity is severe. However, there must also exist a range of temperatures where damage is not additive, and where mechanisms that facilitate recovery can occur. The transition to this range of temperatures is indicated by the sudden break from linearity in the TDT curve of heat stress as shown in Fig. **4C** and in previous studies for plants (e.g. Sapper, 1935; Lorenz, 1939; Levitt, 1980; Nishiyama, 1975; Hara, 2005 for heat stress and Levitt, 1980 for cold stress) and ectotherm animals (e.g. Crozier, 1925; Brown & Crozier, 1927; Jørgensen et al., 2021; Ørsted et al., 2022 for heat stress). The break from linearity denotes a transition from a high temperature range where damage is additive to a lower temperature range where recovery and resistance against thermal stress is possible (Alexandrov, 1964; Kappen and Zeidler, 1977; Ørsted et al., 2022). These temperature zones have previously been referred to as zones of injury and resistance (Alexandrov, 1964). Here these zones are referred to as a stressful temperature range reflecting homeostatic disruption and a permissive temperature range reflecting homeostatic regulation, to avoid confusion about effects of thermal acclimation on thermal tolerance and resistance. The infliction point that separates these temperature zones is known as the critical temperature (*T*_c_) and can be viewed as the temperature or range of temperatures where the net difference between homeostatic regulation and disruption is zero (Jørgensen et al., 2021; Ørsted et al., 2022).

### The stressful temperature range

In the stressful temperature range, temperature acts as an exponentially limiting factor, where the severity of the stress dictates the tolerance duration of a given trait (Brown & Crozier, 1927; Fry et al., 1946; Alexandrov, 1964; Jørgensen et al., 2019, 2021). Conversely, in the permissive temperature range, lifespan indirectly depends on temperature due to its impact on metabolic rates (rate-of-the-living hypothesis; Brown et al., 2004; Price et al., 2010). In this range, various processes such as growth, senescence and general metabolic activities exhibit Q_10_ values around 2 (Běhrádek, 1930; Amthor, 1984; Gillooly et al., 2001; Atkins & Tjoelker, 2003; Allen et al., 2005; Dell et al., 2011; Michaletz, 2018; Rezende & Bozinovic, 2019). However, in the TDT curves presented here for the stressful temperature range of heat stress, as well as in previous studies for plants (Brown & Crozier, 1927; Fel’dman & Liutova, 1962; Alexandrov, 1964), and a range of ectotherm animals (Brown & Crozier, 1927; Cossins & Browler, 1987; Fry et al., 1946; Jørgensen et al., 2019, 2022; Kilgour & McCauley, 1986; Rezende et al., 2014) the Q_10_ values are in the hundreds and even thousands. Although this is not necessarily the case for cold stress (Fig. **4D**), this indicates that plants have a large thermal sensitivity in the extreme temperature ranges. For example, the Q_10_ for Thyme was 3,981 and 3.98*10^19^ for heat and cold stress, respectively. Hence, the exposure time to receive a lethal dose within the stressful range of temperatures for this species decrease with ∼ 129 and 9020 % for every 1 °C increase in the stress intensity for heat and cold stress, respectively. This extreme thermal sensitivity underlines that future frequencies and intensities of thermal extremes as projected by future climate change scenarios could greatly impact the mortality and distribution of many plant species.

Additive damage accumulation suggests that, for a given level of damage received (e.g. 100% damage), the same physiological mechanisms are responsible throughout the entire range of stressful temperatures for a specific trait and type of stress. These mechanisms may largely reflect the direct destructive action of thermal stress to cellular components (e.g. denaturation of proteins, cellular membranes and lipids) (Levitt, 1980; Thomashow, 1999; Ruelland & Zachowski, 2010; Narayanan et al., 2016; Tang et al., 2016; Hayes et al., 2021; Zinta et al., 2022). However, secondary mechanisms that damage various cellular and physiological processes (e.g. transport of metabolites, photosynthesis, respiration, membrane permeability and transport channels) as a result of the original damage may be involved as well (Levitt, 1980; Thomashow, 1999; Atkin & Tjoelker, 2003; Larcher, 2004; Wahid et al., 2007; Prasad et al., 2008; Źróbek-Sokolnik, 2012; Buchanan, 2015; Nievola et al., 2017). In this study, the Q_10_ values for Thyme may reflect the rate of thermal damage accumulation to the thylakoid membranes and thus the thermal tolerance of photosynthesis. However, as the TDT curves in this study were generated 16-19 hours following stress exposure, other secondary actions could also play a role. Q_10_ values of cold stress are often lower than Q_10_ values for heat stress in a range of ectotherms (Rezende et al., 2014) as well as the two examples of plants shown here (Fig. **4D**), and as indicated by studies showing that a very gradual accumulation of damage can occur during cold stress (Fracheboud et al., 1999; Georgieva & Lichtenthaler, 2006; Chaves et al., 2015). However, for Thyme the Q_10_ was many orders of magnitudes greater for cold stress compared to heat stress, and instead of a gradual accumulation of damage, all PSII efficiency was abruptly lost within a narrow temperature range (Fig. **2A**). In this case, damage from cold stress may have largely been driven by the formation of intracellular ice (Levitt, 1980).

### The TDT framework in plant physiological ecology

The application of the TDT framework holds several significant implications and should be considered as a strong complementary tool to study plant thermal tolerance. By using the TDT framework it is possible to compare thermal tolerance estimates among species where the same amount of homeostatic failure has been applied (e.g. the temperatures and stress durations resulting in 100, 50 or 20 % damage), even to species that differ considerably in thermal tolerance (Tarapacki et al., 2021) (e.g. Fig. **4A**). This standardization makes comparisons between species and studies more straightforward and provide a more equal basis for comparison. Plant thermal tolerance is influenced not only by the ability to withstand stressful temperatures when the stress occurs, but also by the capacity to recover from stress and subsequently increase tolerance through hardening (Nievola et al., 2017). The important distinction between the permissive and stressful temperature range through the TDT framework could enhance our understanding of how thermal tolerance is governed by the interplay of accumulated damage, recovery, and hardening mechanisms. For example, Alexandrov (1964) and Alexandrov (1956) showed using TDT curves, that acquired heat hardening is possible when damage has been received in the stressful temperature range followed by recovery in the permissive range. This suggests that damage accumulation is a vital que that may increase thermal tolerance through various physiological responses (Lin et al., 1984; Vierling, 1991; Charng et al., 2007; Wang et al., 2012; Lin et al., 2014; Shinozaki et al., 2015 Younghui et al., 2018; Hayes et al., 2021) and consequently that it may be possible to predict when hardening will occur through the TDT framework. By generating TDT curves, such as in this study, the equation for accumulated damage (Eqn. 3 above) can be used to expose whole plants or different types of tissues to temperature profiles (e.g. by measuring leaf temperatures) resulting in equivalent levels of homeostatic failure (**Notes S1**). Subsequently allowing recovery in the permissive range can illuminate how recovery and hardening processes vary among species at equivalent levels of species-specific homeostatic failures under various environmental conditions over time. This strategic use of TDT curves in experimental designs facilitates the assessment of regulatory rates of these processes and their interplay in sustaining thermal tolerance. Consequently, it offers a more detailed evaluation of how a specific level of homeostatic failure for a given trait relates to overall fitness and mortality. In a similar way, the TDT model may potentially be a useful tool to quantify and predict the temperature-dependent accumulation of damage that indirectly trigger phenological transitions such as flowering, fruiting and leaf senescence (Bahuguna & Jagadish, 2015; Vitra et al., 2017; Grossman, 2023). Hence, the TDT model could offer mechanistic insight into the adaptive potential of populations, communities, and different genotypes in the context of climate change. Similarly, it could be used to assess the likelihood of invasive species successfully colonizing new areas (Zerebecki & Sorte, 2011) or predict how shifts in plant performance due to temperature stress will impact ecosystem dynamics such as carbon sequestration (Dusenge et al., 2019).

The TDT framework can be used to predict when an organism will succumb to thermal stress even when the temperature fluctuates using the equation for accumulated damage (**Notes S1**). Frameworks that predict mortality while taking thermal acclimation into account have already been applied for wild populations of ectotherm animals (Rezende et al., 2020; Jørgensen et al., 2021; Ørsted et al., 2022, Verberk et al., 2023) and recently for plants (Cook et al., 2023, Preprint; Neuner & Buchner, 2023). Although this study confirms the validity of these frameworks, predictions should be made cautiously considering that thermal damage accumulated on a warm day could potentially be reversed during a cooler night, obscuring predictions for the following day (Mathur & Jajoo, 2014). Additionally, this recovery may induce hardening responses within just a few days (Alexandrov, 1964), which will further obscure predictions. Future research should be aimed at testing whether derived TDT model parameters can accurately predict mortality in the field where the temperature fluctuates across T_c_, by comparing predictions with observed mortality levels while accounting for short- and long-term recovery and hardening mechanisms.

Both static and dynamic ramping assays are used to generate estimates of the upper and/or lower thermal limits of species (O’sullivan et al., 2017; Lancaster & Humphreys, 2020; Geange et al., 2021; Wooliver et al., 2022), for example through thermal performance curves (TPCs) which do not take both the intensity and the duration of the stress into account (Rezende & Bozinovic, 2019; O’sullivan et al., 2017). Without taking exposure duration into account, thermal tolerance estimates may be derived from both the permissive and stressful temperature range. These estimates reflect different processes and physiological states and as a result, comparisons may lead to erroneous conclusions about how thermal tolerance vary across species and methods, or how biological and environmental factors affect thermal tolerance. Even when species exhibit equivalent levels of homeostatic failure at a given temperature, variations in thermal tolerance estimates can significantly differ at other temperatures and exposure durations, depending on the species’ thermal sensitivity (Fig. **4D**). However, if damage accumulates in an additive manner, it is possible to mathematically transform one assay type to another (Fry et al., 1946; Jørgensen et al., 2019, 2021; Kilgour & McCauley, 1986; Kilgour et al., 1985) and thus derive TDT model parameters from both static and dynamic assays (Jørgensen et al., 2021; Kilgour & McCauley, 1986). Hence, the TDT model can unify these approaches and enable comparisons based on TTLs (i.e., CT_!_, Q_10_ and T_c_) that align with specific research questions.

## Supporting information

The following is included as supporting information to this manuscript:

**Fig. S1** *F*_V_/*F*_M_ of *Thymus vulgaris* control samples measured in parallel to the static and additive damage accumulation assays of heat and cold stress.

**Fig. S2** *F*_V_/*F*_M_ of *Thymus vulgaris* samples stressed at either 43°C, 43.5 °C, 44 °C, 44.5 °C, 45.5 °C, 46 °C, 47 °C or 47.5 °C for different durations.

**Fig. S3** The proportion of *Thymus vulgaris* samples that were stressed at either -8 °C, -7.5 °C, -7 °C or –6.5

°C for different durations where *F*_V_/*F*_M_ was above a 50% decrease.

**Fig. S4** *F*_V_/*F*_M_ of *Thymus vulgaris* samples stressed at 44 °C and subsequently transferred to either 44.5 °C, 45 °C, 45.5 °C, 46 °C, 46.5 °C or 47 °C for different durations.

**Fig. S5** The proportion of *Thymus vulgaris* samples that were stressed at -6 °C and subsequently transferred to either -7 °C, -7 °C, -7.5 °C, -7.5 °C, -8 °C or -8 °C for different durations where *F*_V_/*F*_M_ was above a 50% decrease.

**Notes S1**: R code to calculate thermal damage accumulation under varying exposure and temperature conditions using the TDT framework.

## Acknowledgements

We thank Lise Lauridsen for helping with maintaining optimal growth conditions for the plants and logistical challenges during the data collection. We thank Laura Skrubbeltrang Hansen for revising the early versions of the manuscript. This work was funded by a grant from Independent Research Fund Denmark (grant number DFF-1026-00173B) to BKE.

## Author contributions

AHF, BE and MØ planned and designed the research. AHF performed experiments and analysed the data. AHF drafted the first version of the manuscript and all authors contributed to and revised the manuscript.

## Data availability

The data used in this manuscript will be made available on https://figshare.com upon acceptance of the manuscript for publication.

