## Supplementary material for "Application of the thermal death time model in predicting thermal damage accumulation in plants": Notes S1

#Accumulated damage function

```
# Insert slope and intercept from a derived TDT curve.
# Below is the slope and intercept from the TDT curve
# based on heat stress in Thyme.
slope = -0.3597205
intercept = 18.03221

# This function calculates accumulated damage across
# a temperature profile depending on the exposure duration
# according to equation 3.
Acc_function = function(x, y) {
  output = data.frame(z = NA, cumsum = NA)

  # Duplicate the last row of the input data frame
  last_row = nrow(data.frame(x = x, y = y))
  x = c(x, x[last_row])
  y = c(y, y[last_row])

  for (i in 1:length(x) - 1) {
    print(paste0("round nr. ", i))
    acc = max(0, min(100, (100 * (x[i + 1] - x[i])) / (10^(slope * max(y[i + 1], y[i])
                                                            + intercept))))
    output[i, 1] = acc
  }

  output$cumsum = cumsum(output$z)
  z = output$cumsum
  z = append(z, 0)
  z[z > 100] = 100  # Set values exceeding 100% to 100%

  # Remove the last row of the calculations
  z = z[-length(z)]

  return(z)
}
```

#Temperature profile

```
# This code creates a fluctuating temperature and exposure duration profile.
# In this illustrative example, the temperature fluctuates between
# 40 to 47 degrees celsius, and the plant is exposed to each temperature
# for 10 minutes.
specific_temperatures <- c(43, 43.5, 44, 44.5, 45, 44, 43, 44, 45, 46, 47)
```

```r
Minutes_at_each_temp <- 10

Temperature <- rep(specific_temperatures, each = Minutes_at_each_temp)
Time <- rep(seq(1, Minutes_at_each_temp), times = length(specific_temperatures))

data <- data.frame(Time,Temperature)
data$Time <- seq(1, nrow(data))

# Here the temperatures and exposure durations are inserted in the accumulated damage
# function. The function calculates accumulated damage across the temperatures and
# exposure durations. Any profile of temperatures and exposure durations can be
# passed to this function.
data$Acc_damage = Acc_function(data$Time,data$Temperature)
```

#Plotting the result #Temperature profile

```r
library(ggplot2)
data$Time <- as.numeric(data$Time)
data$Temperature <- as.numeric(data$Temperature)
data$Acc_damage  <- as.numeric(data$Acc_damage)
legend_title <- "Damage (%)"
colfunc<-colorRampPalette(c("honeydew","gray21"))

TDT_curve <- ggplot(data, aes(x = Time, y = Temperature)) +

    geom_point(data = data, aes(x = Time, y = Temperature, fill = Acc_damage),
               shape = 21, size = 3,alpha = 1) +

 scale_fill_gradientn(legend_title,colours = colfunc(21)) +
 geom_hline(yintercept = 100, linetype = "dashed") +

  labs(x= "Time (minutes)", y = expression(paste("Temperature"," (\u00B0C)"))) +

theme(legend.title = element_text(color = "black", size = 16, family = "Times",
face = "bold"), legend.text = element_text(color = "black", size = 16,
                                           family = "Times"),
legend.position = "right",
legend.key = element_rect(color = "transparent", fill = "transparent"),
legend.background = element_rect(fill="transparent", size=0.5,
                                 colour ="transparent"),
  # Remove panel border
  panel.border = element_blank(),
  axis.ticks.length=unit(.25, "cm"),
  # Remove panel grid lines
  panel.grid.major = element_blank(),
  panel.grid.minor = element_blank(),
  # Remove panel background
  panel.background = element_blank(),
  # Adjust axis
  axis.line = element_line(colour = "black"),
  axis.text.x = element_text(size = 20, colour = "black", family = "Times"),
  axis.text.y = element_text(size = 20, colour = "black", family = "Times"),
  axis.title.x = element_text(size = 20, colour = "black", family = "Times"),
```

```
  axis.title.y = element_text(size = 20,
  colour = "black", family = "Times")) + scale_x_continuous(breaks
 = c(0,20,40,60,80,100,120,140,160)) +
  coord_cartesian(xlim =c(0, length(data$Time)),
  ylim = c(42.5,47.5)) + scale_y_continuous(breaks = c(43,44,45,46,47))
print(TDT_curve)
```

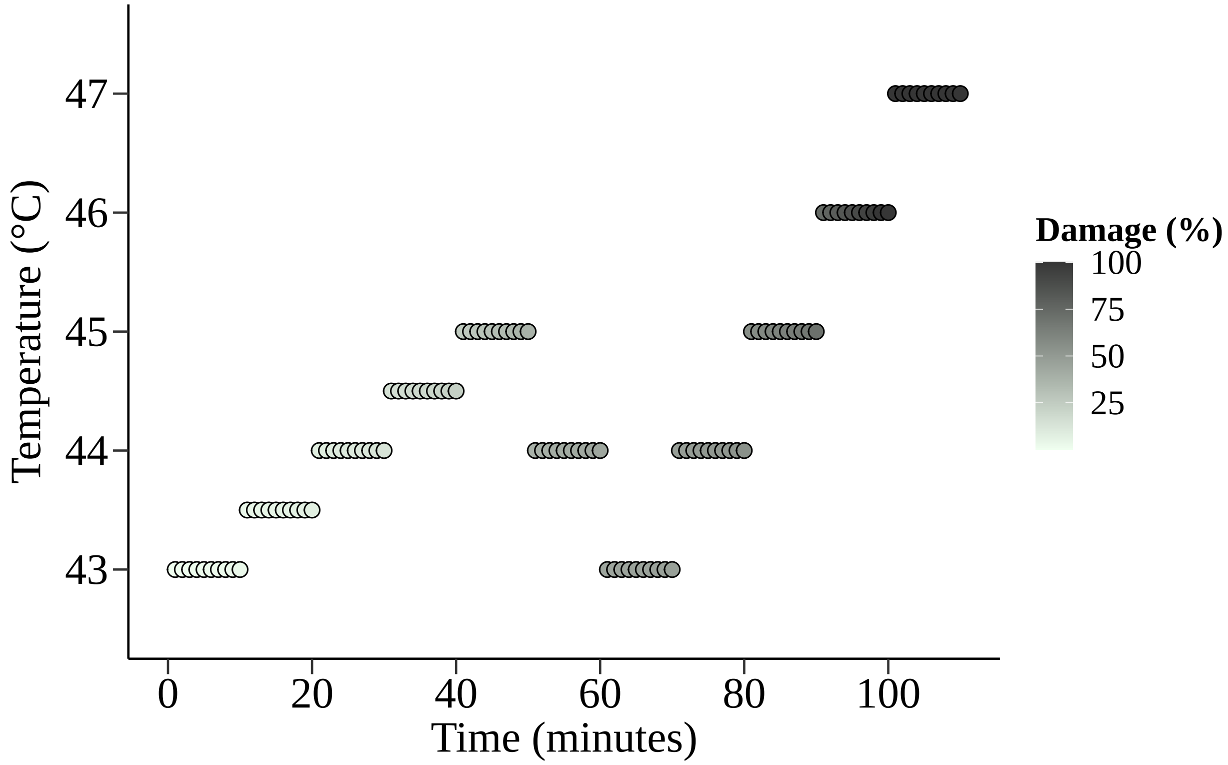


#Accumulated damage

```
library(ggplot2)
data$Time <- as.numeric(data$Time)
data$Temperature <- as.numeric(data$Temperature)
data$Acc_damage  <- as.numeric(data$Acc_damage)
legend_title <- "Temperature (\u00B0C)"
colfunc<-colorRampPalette(c("honeydew","gray21"))


TDT_curve <- ggplot(data, aes(x = Time, y = Acc_damage)) +

    geom_point(data = data, aes(x = Time, y = Acc_damage, fill = Temperature)
               ,shape = 21, size = 3,alpha = 1) +

 scale_fill_gradientn(legend_title,colours = colfunc(21)) +
 geom_hline(yintercept = 100, linetype = "dashed") +

  labs(x= "Time (minutes)", y = expression(paste("Accumulated damage"," (%)"))) +

theme(legend.title = element_text(color = "black", size = 16,
```

```
              family = "Times", face = "bold"),
      legend.text = element_text(color = "black",
                                 size = 16, family = "Times"),
      legend.position = "right",
legend.key = element_rect(color = "transparent", fill = "transparent"),
legend.background = element_rect(fill="transparent", size=0.5,
                                 colour ="transparent"),
  # Remove panel border
  panel.border = element_blank(),
  axis.ticks.length=unit(.25, "cm"),
  # Remove panel grid lines
  panel.grid.major = element_blank(),
  panel.grid.minor = element_blank(),
  # Remove panel background
  panel.background = element_blank(),
  # Adjust axis
  axis.line = element_line(colour = "black"),
  axis.text.x = element_text(size = 20, colour = "black", family = "Times"),
  axis.text.y = element_text(size = 20, colour = "black", family = "Times"),
  axis.title.x = element_text(size = 20, colour = "black", family = "Times"),
  axis.title.y = element_text(size = 20,
  colour = "black", family = "Times")) + scale_x_continuous(
    breaks = c(0,20,40,60,80,100,120,140,160)) +
  coord_cartesian(xlim =c(0, length(data$Time)),
  ylim = c(0,110)) + scale_y_continuous(breaks = c(0,20,40,60,80,100))
print(TDT_curve)
```

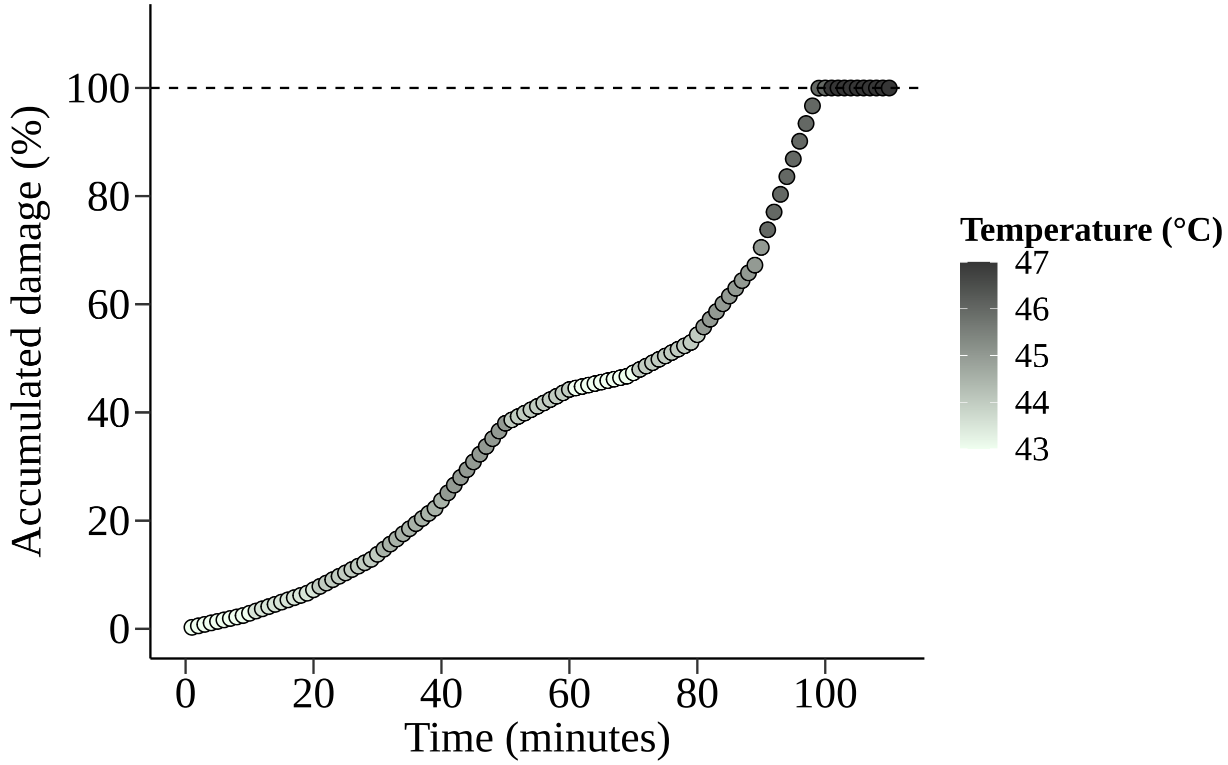
