## Supplementary material for "Application of the thermal death time model in predicting thermal damage accumulation in plants": Fig. S1

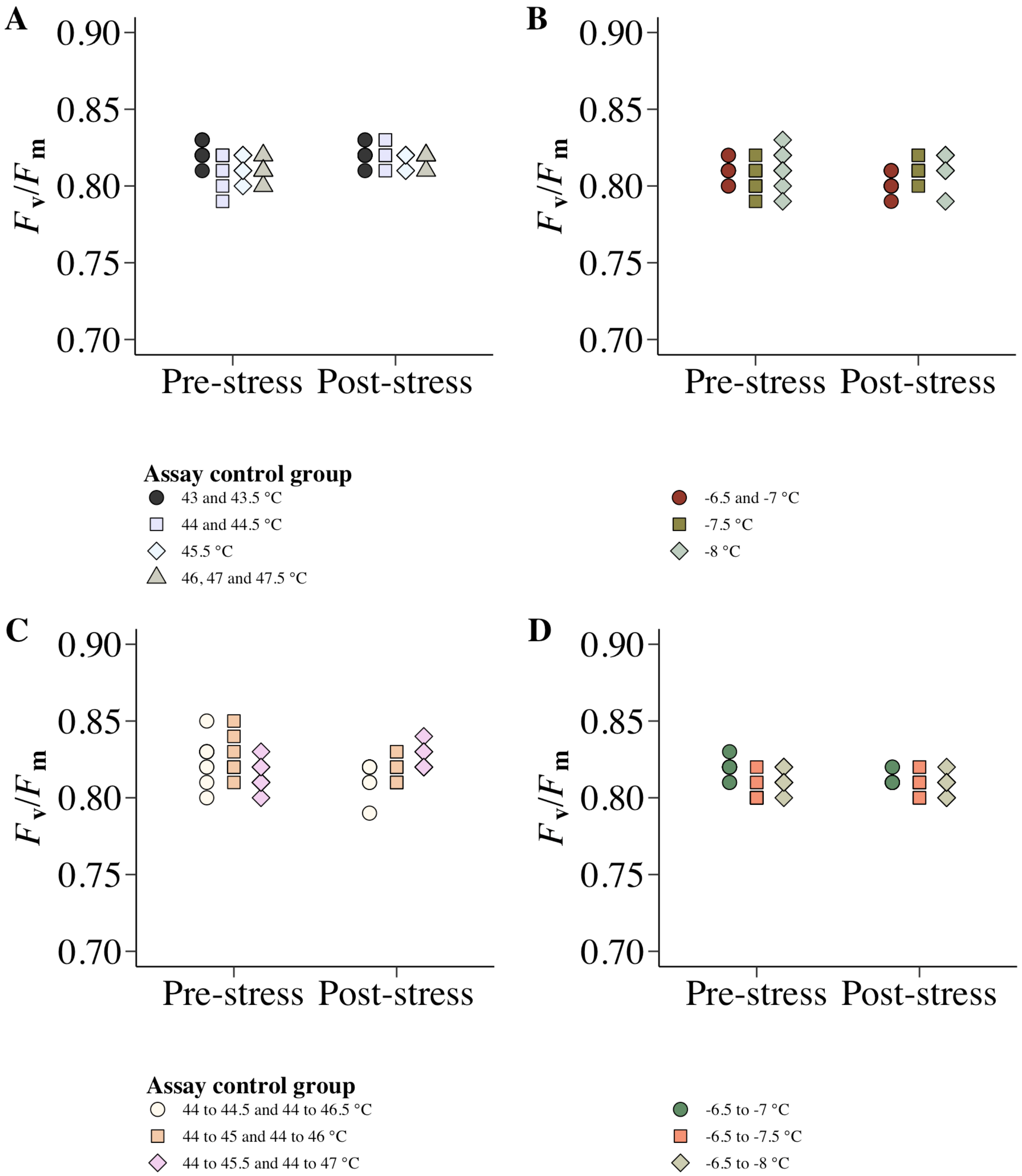


**Fig. S1** $F_V/F_M$ of *Thymus vulgaris* control samples measured in parallel to the **A-B)** static and **C-D)** additive damage accumulation assays of heat and cold stress. Different symbols indicate the stress treatments that the controls were measured in parallel to (n=10 for each treatment). The control samples were used to confirm that $F_V/F_M$ was identical before and after the experimental period when the samples were not stressed.
