## Supplementary material for "Application of the thermal death time model in predicting thermal damage accumulation in plants": Fig. S2

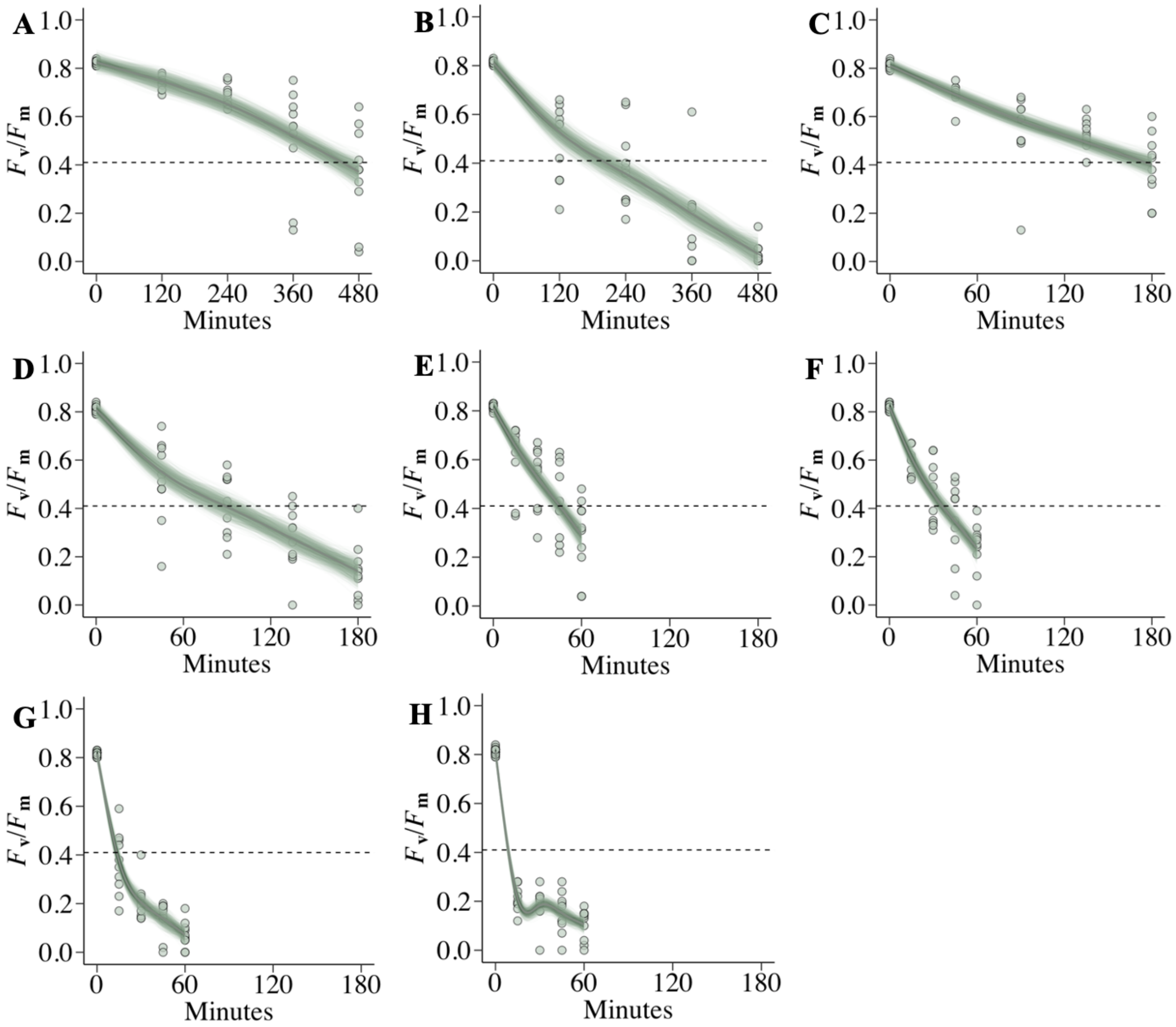


**Fig. S2** $F_V/F_M$ of *Thymus vulgaris* samples stressed at either **A)** 43°C, **B)** 43.5 °C, **C)** 44 °C, **D)** 44.5 °C, **E)** 45.5 °C, **F)** 46 °C, **G)** 47 °C or **H)** 47.5 °C for different durations (n=40 for each treatment). Generalized additive models were fitted to the time response of $F_V/F_M$ for each treatment with 10.000 random draws from the multivariate normal distribution of the model parameters. The time for $F_V/F_M$ to decrease with 50% is indicated when the fitted GAMs cross the dotted line representing $CT_\tau$. $CT_\tau$ denotes the critical temperature at which a lethal dose of damage to PSII occurs following a specific time of exposure. $F_V/F_M$ of all samples were initially measured prior the stress and are presented at the exposure time of zero minutes.
