## Supplementary material for "Application of the thermal death time model in predicting thermal damage accumulation in plants": Fig. S3

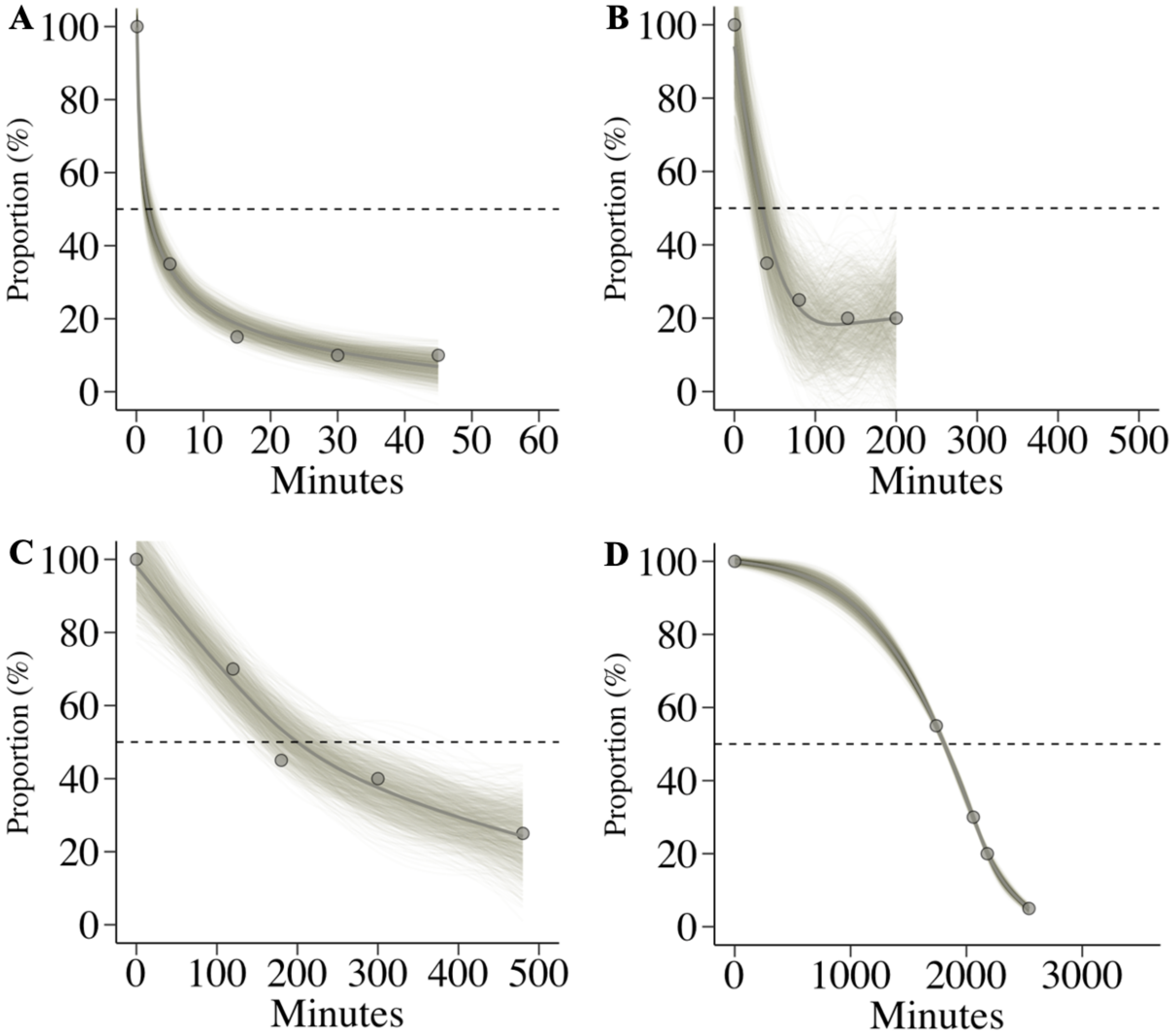


**Fig. S3** The proportion of *Thymus vulgaris* samples that were stressed at either **A)** -8 °C, **B)** -7.5 °C, **C)** -7 °C or **D)** –6.5 °C for different durations where $F_V/F_M$ was above a 50% decrease. Generalized additive models were fitted to the time response of the proportion of samples where $F_V/F_M$ was above a 50% decrease. The models were fitted with 10.000 random draws from the multivariate normal distribution of the model parameters. The time when 50% of samples had more than a 50% decrease in $F_V/F_M$ is indicated when the fitted GAMs cross the dotted line representing $CT_\tau$. $CT_\tau$ denotes the critical temperature at which a lethal dose of damage to PSII occurs following a specific time of exposure. Individual symbols represent 20 samples and $F_V/F_M$ of all samples was initially measured prior the stress which is presented at the exposure time of zero minutes.
