## Supplementary material for "Application of the thermal death time model in predicting thermal damage accumulation in plants": Fig. S4

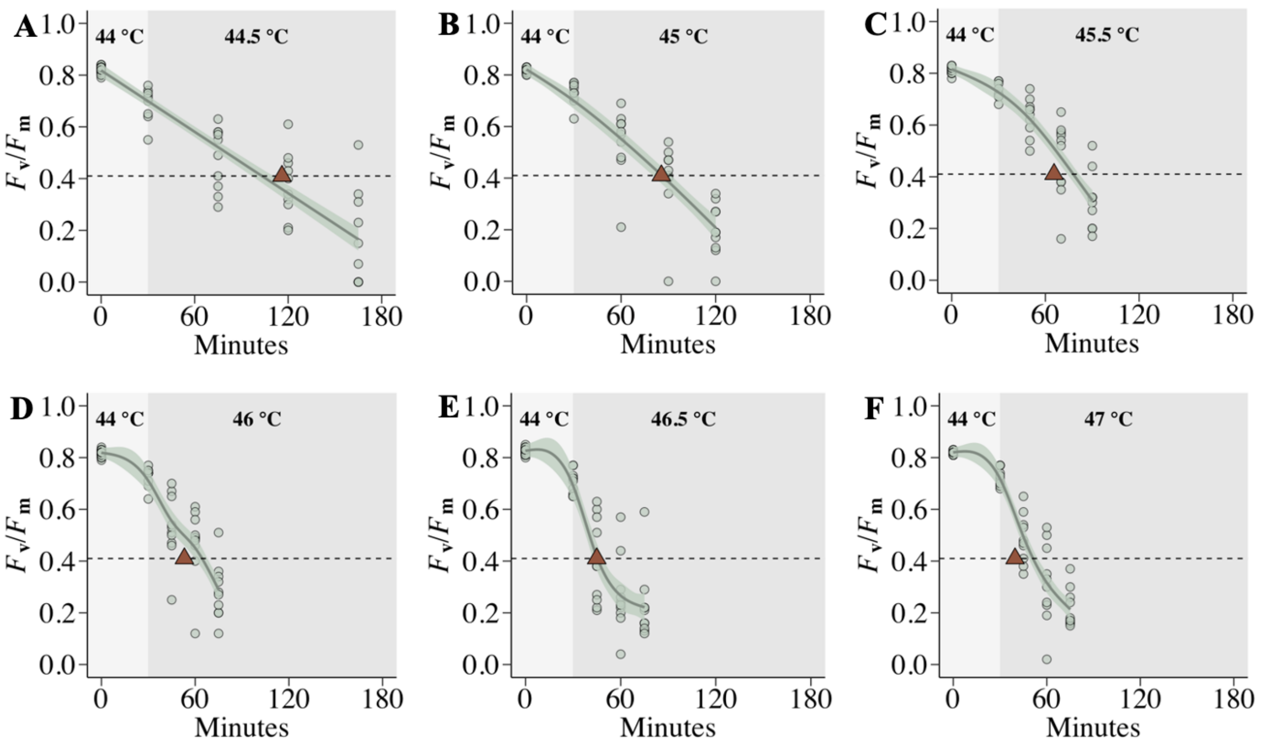


**Fig. S4** $F_V/F_M$ of *Thymus vulgaris* samples stressed at 44 °C (denoted by the light shaded areas) and subsequently transferred to either **A)** 44.5 °C, **B)** 45 °C, **C)** 45.5 °C, **D)** 46 °C, **E)** 46.5 °C or **F)** 47 °C (denoted by the dark shaded areas) for different durations (n=40 for each treatment). Generalized additive models were fitted to the time response of $F_V/F_M$ for each treatment with 95% point-wise confidence intervals. The time for $F_V/F_M$ to decrease with 50% is indicated when the fitted GAMs cross the dotted line representing $CT_\tau$. $CT_\tau$ denotes the critical temperature at which a lethal dose of damage to PSII occurs following a specific time of exposure. $F_V/F_M$ of all samples were initially measured prior the stress and are presented at the exposure time of zero minutes. Thermal death time curves were used to predict $CT_\tau$ for each temperature treatment, and the predictions are denoted by the red triangles. Individual samples are denoted by the closed circles.
