## Supplementary material for "Application of the thermal death time model in predicting thermal damage accumulation in plants": Fig. S5

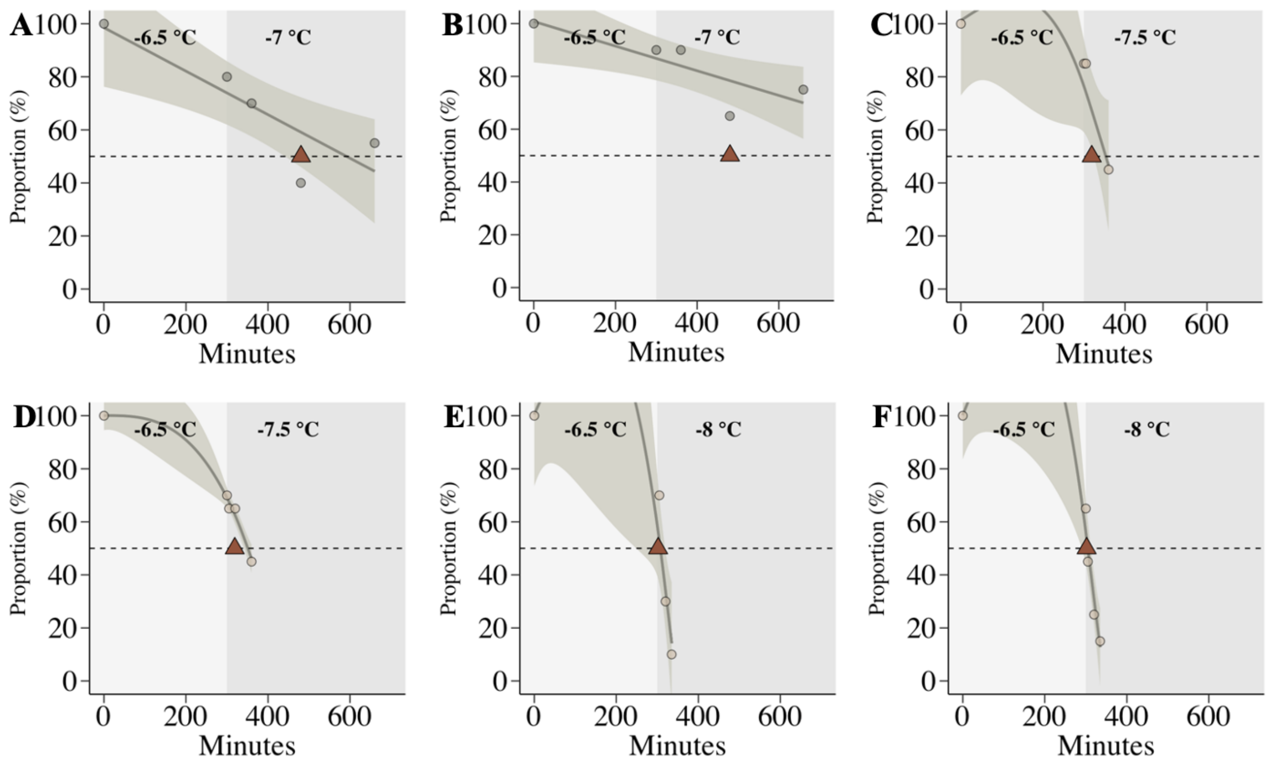


**Fig. S5** The proportion of *Thymus vulgaris* samples that were stressed at -6 °C (denoted by the light shaded areas) and subsequently transferred to either **A)** -7 °C, **B)** -7 °C, **C)** -7.5 °C, **D)** -7.5 °C, **E)** -8 °C or **F)** -8 °C (denoted by the dark shaded areas) for different durations where $F_V/F_M$ was above a 50% decrease. Generalized additive models were fitted to the time response of the proportion of samples where $F_V/F_M$ was above a 50% decrease with 95% point-wise confidence intervals. The time when 50% of samples had more than a 50% decrease in $F_V/F_M$ is indicated when the fitted GAMs cross the dotted line representing $CT_\tau$. $CT_\tau$ denotes the critical temperature at which a lethal dose of damage to PSII occurs following a specific time of exposure. Thermal death time curves were used to predict $CT_\tau$ for each temperature treatment, and the predictions are denoted by the red triangles. Closed circles represent 20 samples and $F_V/F_M$ of all samples was initially measured prior the stress which is presented at the exposure time of zero minutes.
